# Toward Electrical Impedance Tomography of Functional Activity in the Retina

**DOI:** 10.64898/2026.10.23.761326

**Authors:** Enrico Ravagli, Alistair McEwan, Dario A. Protti, Kirill Aristovich

**Affiliations:** Department of Medical Physics and Biomedical Engineering, University College London, London, United Kingdom; School of Medicine and Population Health, University of Sheffield, UK; Department of Medical Physics and Biomedical Engineering, University College London, UK; School of Medical Sciences, University of Sydney, Sydney, Australia; School of Biomedical Engineering, University of Sydney, Sydney, Australia

**Keywords:** Retina, Multi-Electrode Array, MEA, Electrical Impedance Tomography, EIT, Vision, Prosthetics, Electrophysiology, Neuroscience

## Abstract

Electrical Impedance Tomography (EIT) is a non-invasive electrical imaging technique that reconstructs spatial variations in tissue conductivity. Traditionally used for thoracic imaging, EIT has recently been extended to neural applications in the brain and peripheral nerves. This study investigates the feasibility of using EIT to image functional neural activity in the retina. We present computational modeling of EIT measurements for image reconstruction of small conductivity perturbations across different layers of the ex-vivo mouse retina with electrodes placed in epiretinal configuration. Experimentally, we describe the design of a multi-electrode array suitable for impedance measurements under these conditions and report preliminary recordings of light-evoked impedance changes in response to wide-field optical retinal stimulation. Our computational results suggest that small conductivity perturbations of 200µm diameter and 5% conductivity variation are experimentally detectable by high-end biopotential recording equipment and could be localized by EIT imaging within ≈13% of inter-electrode distance and ≈22% of retinal thickness, respectively. In preliminary recordings, we observed slow impedance variations of 13.0±9.3 µV (0.37±0.44%) in response to retina-wide light pulse trains, when performing impedance measurements with 10 µA applied current at 2-8 kHz. These variations were characterized by a slow time course, distinct from the fast neural EIT signals associated with action potentials, and may reflect extracellular or metabolic processes linked to retinal activation. Our modelling and technical results support the feasibility of retinal EIT, while the experimental observations are preliminary and motivate further controlled studies. Retinal EIT could significantly advance retinal electrophysiology research and prosthetic development.

## I. Introduction

Electrical impedance tomography (EIT) is a non-invasive technique which allows imaging of variations in electrical impedance inside a volume of interest by image reconstruction of transfer impedance measurements collected from several pairs of external electrodes [1]. This technique features a much lower spatial resolution compared to other types of tomographic techniques such as Magnetic Resonance Imaging (MRI) or X-Ray Computed Tomography (CT), but it allows for a much higher temporal resolution, in the order of milliseconds, and might be performed with compact and relatively inexpensive hardware. As such, EIT can provide continuous or semi-continuous monitoring. More so, EIT hardware is purely electronic and lends itself easily to miniaturization for wearable or implantable applications [2].

While historically and traditionally applied to thoracic imaging, EIT has more recently been proposed for several types of biological tissues and biomedical applications, such as imaging for breast or abdomen [3]. One particular branch of novel EIT applications involves imaging of neural tissue; within the last 10 years, EIT has been applied to imaging physiologically- or electrically-evoked compound neural activity in brain and nerves [4], [5], [6], [7] and phenomena such as stroke and epileptic events [8], [9], [10]. In the former case, labelled as “fast neural EIT”, neuronal activity is imaged by the detection of small variations in electrical impedance produced by the opening of ion channels during action potentials, and requires averaging to reach a Signal-to-noise Ratio (SNR) sufficient for imaging. In the latter case, slower, single-shot events such as an osmotic water shift or blood haemorrhage are detected by the technique. A distinguishing feature for these different sources of impedance variation is their time course; fast neural changes occur on the order of milliseconds (≈1 kHz), whereas changes linked to extracellular phenomena evolve on sub-second timescales (<1 Hz).

The purpose of this study was to investigate the feasibility of EIT as an imaging technique for detecting functional activity in the retina.

The retina is the biological structure of the eye responsible for the process of photo-electrical transduction that results in visual inputs encoded for transmission through the optic nerve to different visual areas in the brain [11]. In the healthy retina, this process is performed by a complex layered neural structure; light impacting photoreceptor cells is converted to electrical signals that travel multiple layers to reach the most external one made up of retinal ganglion cells, while being subject to processing for feature extraction on its way. However, in a number of degenerative retinal diseases, visual perception is impeded as a consequence of loss of photoreceptor cells.

Examples of such diseases are Retinitis Pigmentosa (RP) and Age-related Macular Disease (AMD), which are leading causes of irreversible vision loss. The worldwide prevalence of RP is about 1 in 4000, for a total of more than 1 million affected individuals [12], while AMD affects more than 8.7% of the population worldwide[13]. Despite loss of function of photoreceptor cells in these diseases, other layers of retinal neurons are generally preserved [14]; retinal prosthesis systems aim to take advantage of this fact by inducing perception of visual effects through electrical stimulation of the retina with implantable devices [15].

In the last 25 years, retinal prostheses have undergone significant advances, with multiple engineering approaches tested in terms of materials, electronics, signal processing and surgical procedures [16]. One critical aspect that remains a strong focus of current research is the identification of optimal spatial and temporal patterns of electrical stimulation needed to elicit retinal activity that will lead to visual perception [17]. While modelling and experiments [18] support validation of such patterns, the development of a non-invasive technique for imaging the functional neural activity of the retina, either as a result of light stimulation in a healthy retina or electrical stimulation from a retinal prosthetic in a diseased retina, would greatly accelerate progress in this field. Another recent line of research for the treatment of retinal degenerative diseases is the use of optogenetic techniques for vision restoration [19]. This group of techniques typically involves expression of light-sensitive proteins in layers of the retina different from the one subject to degeneration, such as the retinal ganglion cell (RGC) layer. However, this approach presents distinct challenges, including the need to precisely identify and target the appropriate retinal site for delivery of viral vectors encoding the photosensitive proteins[20].

EIT represents an innovative tool which potentially could perform imaging of retinal functional activity and provide spatial information about retinal processing of incoming optical and electrical stimuli. This could support both basic electrophysiology research, optogenetic research, and retinal prosthetic design and operation. To the best of our knowledge, only very limited work has been performed in the past regarding bioimpedance and EIT applied to retina. In 1970, Ogden and Ito analysed the resistivity of avian retinae with a particular interest in tissue anisotropy. In 1982, Ferreira-Filho and Martins-Ferreira [22] measured the electrical impedance of isolated chick and toad retinae in the absence and presence of slow waves of suppressed neuronal activity. In 1985, Karwoski et al. conducted experiments to record the laminar profile of resistivity in frog retina [23]. In more recent times, other groups also reported studies focused on analysis of retinal resistivity [24], [25].

With this study, we aimed to:

i. Demonstrate proof-of-concept of EIT use for retinal imaging, by performing modelling and experiments in a simplified experimental context and detect impedance changes in response to light stimuli
ii. Provide the scientific community with a set of technical recommendations as a starting point for further retinal EIT research

## II. Study Design and Outline

To investigate the basic feasibility of retinal EIT, we focused our study on ex-vivo impedance measurements of perfused mouse retinas mounted on a planar Multi-Electrode Array (MEA), while applying light pulses to evoke functional activity. Recordings were initially made using various combinations of injected EIT current frequencies and with light stimuli applied at different intensities, wavelengths, pulse durations, and repetition rates; as the study progressed, we focused on a specific pattern of optical stimulation consisting of bursts of light pulses.

Initially, commercially available MEAs were used for recordings, but these proved incompatible for long-term use with our existing neural EIT recording system [26]. As a result, we switched to custom-made MEAs. A detailed discussion of this technical issue can be found in Supplement Section S1. The creation of custom MEAs involved evaluating various technologies based on factors such as ease of manufacture, suitability for our project, electrode electrical properties, and resistance to photoelectric artifacts.

Experimental recordings were preceded by, and conducted in parallel with, computational modeling. The modeling aimed to identify an optimal EIT current injection pattern to be used in experiments, that would maximize current density distribution across the ex-vivo retina, thereby enhancing imaging resolution. Once the ideal EIT measurement pattern was determined, we also simulated small conductivity variations at different spatial locations within the retina and fed these into our reconstruction algorithm to estimate expected impedance changes. Impedance changes expected from modeling were then compared to the ones collected experimentally.

## III. Materials and Methods

### A. Modeling, EIT protocol optimization, and Image Reconstruction

Modeling was performed according to our previously published workflow [6], [7]. Briefly, a Finite-Element (FE) model was developed which represents a retina placed inside a perfusion chamber (Fig.1). Electrodes were placed on the model boundary at the bottom to represent contact between the retina and MEA. Size, number and spacing of simulated electrodes were in accordance with the geometry of our custom FlexPCB array.

**Fig. 1.**
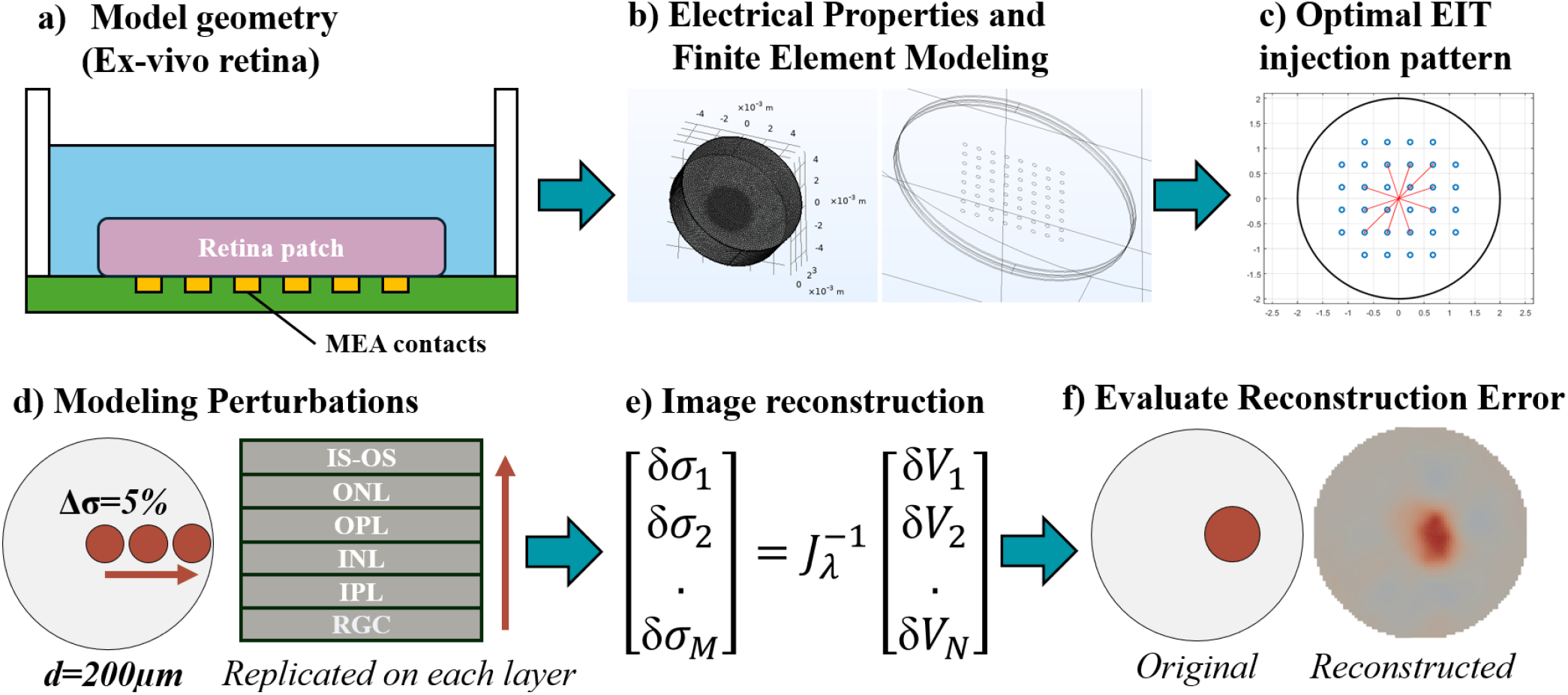
Modeling Workflow – a) A conceptual model is established based on geometry of the problem. b) A tetrahedral mesh is generated including retinal layers, external perfusion fluid, and the position of electrodes, to run FEM simulations. c) An optimization algorithm is applied to find the EIT current injection protocol which maximizes current density in the retina. d) Simulated data is obtained from the same model by applying small conductivity changes. e) Image reconstruction is performed on a coarse version of the mesh, and f) accuracy of reconstructed images is assessed quantitatively.

The retinal domain was represented by a cylindrical domain of 3.5mm diameter and 0.185mm height, fully contained inside another cylindrical domain of 10mm diameter and 3mm height, representing perfusion fluid. The retinal domain was further divided into subdomains, stacked vertically, representing different physio-anatomical layers of the mouse retina from the inner/outer segment to the ganglion cells layer.

Thickness and electrical conductivity of each layer were assigned based on experimental data for mouse retina whenever available, and on data from other species (e.g. rat, human) otherwise [27], [28], [29], [30], [31], [32]. Chosen values are reported in Table 1. Electrical conductivity of the perfusion fluid domain was set to 1.5 S/m.

**TABLE I.** Properties of Retinal FEM Subdomains.

| Layer | Thickness [$\mu\text{m}$] | Conductivity [S/m] |
| --- | --- | --- |
| RGC | 20 | 1.00 |
| IPL | 45 | 0.41 |
| INL | 27 | 0.27 |
| OPL | 15 | 0.29 |
| ONL | 55 | 0.50 |
| IS-OS | 23 | 0.7 |

RGC: Retinal Ganglion Cells; IPL: Inner Plexiform Layer; INL: Inner Nuclear Layer; OPL: Outer Plexiform Layer; ONL: Outer Nuclear Layer; IS-OS: Inner/Outer Segment.

The model was converted to a 7.8M-elements tetrahedral mesh for the purpose of computing the forward problem solution, using the complete electrode model (CEM) within the EIDORS MATLAB plug-in [33]. Electrode contact impedance in the model was set to 2kΩ, compatibly with experimental values assessed by Electrica Impedance Spectroscopy (EIS).

A method previously developed in our group [28] was employed to identify the optimal electrode addressing protocol, aimed at maximizing current density in a region of interest, which in this study corresponded to the entire retina. A total of six current injections, corresponding to six independent MEA electrode pairs, were selected to balance the amount of collected data against the recording duration.

The FEM model was also used for EIT simulations and image reconstruction. Simulated EIT recordings were generated by applying a localized 5% conductivity increase over spots with 200µm diameter, spaced at 0-1100 µm from the center of the retina in 100µm steps, on each of the six retinal layers, for a total of 72 perturbations. Noise was added to the simulated recordings by random number generation, with amplitudes comparable to experimental values (**≈** 0.1 µV).

EIT image reconstruction from simulated and experimentally collected data was performed using noise-corrected 0^th^ order Tikhonov regularization, as done in previous work by our group [6], [7]. The original 7.8M tetrahedral FEM mesh was converted to a 31k-elements hexahedral voxel-based mesh, for the purpose of facilitating image visualization and reducing degrees of freedom during Jacobian matrix inversion. Voxel size was set to 40×40×40µm.

Locations of reconstructed conductivity perturbations within the retina were compared to the location of the corresponding simulated perturbations by computing Centre-of-Mass (CoM) error over the radial and vertical axes.

### B. Multi-Electrode Arrays

Preliminary experiments in this study were performed using a commercially available MEA (60PEDOT-pMEA200/30iR-Ti-Au, Multi Channel Systems, Reutlingen, Germany). After the first few experiments, this array was replaced by a custom MEA due to the high electrode impedance (≈20kΩ), not ideal for EIT recordings, and also due to systematic signal drift observed when recording with our neural EIT system. Details regarding our signal drift issue are reported in Supplement Section S1.

These issues with the standard commercial MEA led us to design our own custom MEA. We manufactured a small number of versions based on traditional rigid PCB technology, and coated electrodes with PEDOT:pTS conductive polymer to decrease the phase and magnitude of contact impedance, as done in previous studies by our group [34]. However, in a limited early set of experiments we observed substantial photoelectrical artefacts in the light-evoked retinal recordings obtained with this combination of materials and led us to our final MEA design. Please see Supplement Section S2 for more details on this photosensitivity issue.

The final version of our custom MEA (Flex32, Fig.4a-b) is based on a Flexible PCB circuit: in this design, 32 electrodes of 100µm diameter are arranged in a 6×6 grid with missing corners and spaced at 450µm intervals. Connecting tracks and electrode are made of copper with 18 µm thickness. Exposed electrode area was ENIG-coated by the PCB manufacturer with a 3µIn thickness. To further reduce phase and magnitude of contact impedance when interfaced with the tissue, electrodes were also subject to platinum coating. Coating was performed manually by applying platinum solution (Gold Solution Plating, UK) with an electroplating pen (SPA Plating, UK) applying a voltage of 8V. To allow for suction, holes of 100µm diameter were drilled in the space in-between electrodes.

Both the commercial (MCS) and custom (Flex32) MEAs feature a rigid chamber glued on top of the MEA to allow for perfusion fluid to supply the *ex-vivo* tissue. In the case of the custom MEA, an acrylic disk was glued to the MEA with silicone of 21mm internal diameter and 2mm thickness was attached to the MEA post-coating with silicone glue. Both the MCS and FlexPCB MEAs were designed to be compatible with the standardized MEA60 interface.

EIS was performed in the 2-10kHz frequency range on a subset of electrodes while the Flex32 was filled with physiological saline solution (NaCl 0.9% w/v) to assess magnitude and phase of contact impedance.

Electrical recordings from MEAs during changing lighting conditions, such as in the case of light pulses applied in the present study, are prone to photoelectric artefacts [35]. We assessed the risk of photoelectric signal artefacts in our newly developed MEA by applying light pulses of medium-high intensity and observing signal response in experimentally relevant bandwidths.

A preliminary test of basic functionality for EIT imaging was performed by 3D-printing a mechanical support and a small thin cylindrical perturbation and dropping the perturbation in the center of the MEA. EIT was performed with 10µA current at 6kHz frequency in time-difference mode (no perturbation vs presence of perturbation).

### C. Surgical Procedures and Ex-vivo Tissue Handling

Animal experiments were performed at the University of Sydney, NSW, Australia. This study followed the NSW *Animal Research Act 1985* and was authorized by the local Animal Research Authority (Ethical Approval 2024/2411).

The study was performed using male C57/BL6J mice, all within 6-12 weeks of age, and experiments were performed within 8 weeks after receipt. On the day of the experiment, mice were transported to the lab and placed in dedicated housing for dark adaptation, which lasted 1.5-2 hours. Briefly, mice were euthanized, and their eyes were enucleated. The cornea, iris and vitreous humor were removed from the eye and the retina separated from the underlying pigment epithelium and sclera. Excised retinae were placed in a 3-D printed holding device and then transferred to our experimental setup. The tissue was continuously perfused with carboxygenated AMES medium at 3–5 mL/min at 35^°^C.

### D. Experimental Setup and Recording Protocols

Experiments were performed in a dedicated retinal electrophysiology setup (Fig. 2).

**Fig. 2.**
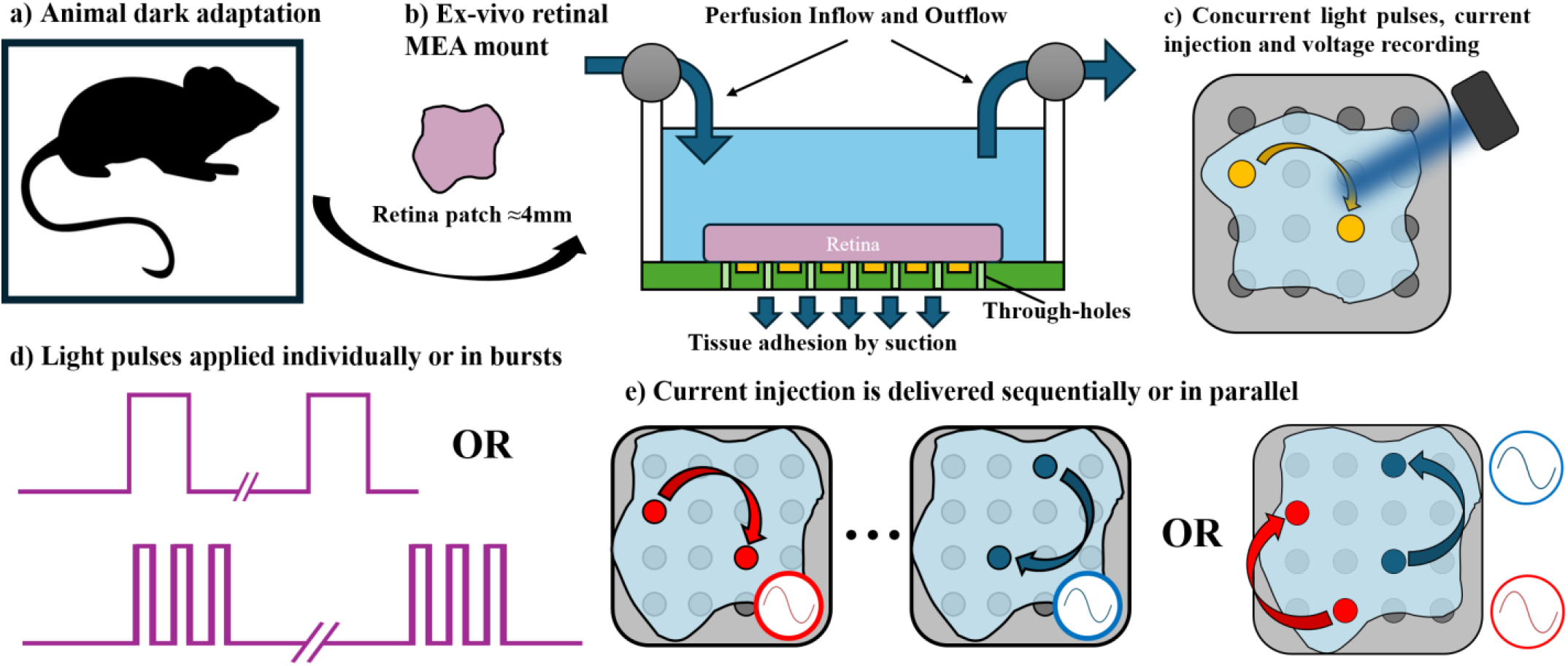
Experimental Workflow – a) Mice was dark-adapted for 2-hours. b) A patch of retinal tissue was mounted on an MEA and perfused. c) Light pulses were delivered to the tissue while injecting electrical current and recording voltage on each electrode. d) Light pulses were either individual or bursts. e) Current injection to electrode pairs was delivered either serially or in parallel.

The setup was composed of a PPS2 peristaltic perfusion system (Multi Channel Systems, Reutlingen, Germany), an upright microscope (BX50 WI, Olympus Corp., Tokyo, Japan), a full-field LED light source (DC4100, Thorlabs, New Jersey, USA) with multiple central wavelengths, an optical microscope and digital camera, and a chamber which allowed interconnection between the MEA and an external recording device. The setup was enclosed into a metal mesh connected to mains ground to decrease the effect of external electromagnetic noise and interference.

Spontaneous and evoked electrophysiological activity, and EIT recordings, were performed using the Actichamp biopotential recorder (Brain Products GmbH, Gilching, Germany) connected to the MEA though a custom adapter PCB. Recordings were performed using a small Ag-AgCl pellet, placed inside the MEA chamber, as the device’s physical ground, and one of the MEA electrodes as the digital reference.

For EIT recordings, sinusoidal current was driven through selected electrode pairs using either a single-channel[36] or multi-channel [37] custom device based on Enhanced Howland Current Source (EHCS) topology. Earlier experiments in the study were performed with the single-channel current source using current with 10µA amplitude and a frequency range of 2-8 kHz. Further into the study, we replaced sequential injection with parallel-mode EIT, injecting current in the amplitude range of 5-10µA; in this modality, all 6 available channels were used simultaneously, with injection frequencies equally spaced at either 1kHz steps in the 3-8khz range, or 100Hz steps in the ranges of 3.2-3.7 kHz, 5.2-5.7 kHz, 7.2-7.7 kHz.

Light pulses to evoke neural activity were delivered as full-field illumination in either individual pulses or pulse trains. For individual pulses, pulse duration was set to 200ms, and repetition rate was 5s. For pulse trains, groups of 20pulses of 200ms on/off duration were evoked with 5-10s in between bursts. Individual light pulses or bursts were repeated 5 times for each recording and averaged for improving SNR of recordings. For this study we used a blue light source with a central optical wavelength of 465nm. Optical pulse intensity was in the range of 14.5-15.5 log photons·cm^−2^·s^−1^. Digital control of the light source and neutral density (ND) filters were used to modulate light intensity and adjust stimulus to the response of individual retinae.

A more detailed report on experimental recordings, including a list of settings selected for each experiment can be found in Supplement Section S5. Electrophysiological control recordings and data processing were performed and are reported in Supplement Section S6.

### E. Signal Processing and Data Analysis

Recordings of spontaneous activity and light-evoked responses were subject to band-pass filtering to extract either the compound microElectroRetinoGram signal (MicroERG) or spiking signals indicative of detection of localized single-unit or multi-unit activity. For MicroERG recordings, a digital 1-300 Hz bandpass filter was first applied, followed by a visual inspection to identify the MEA channels with the strongest MicroERG signals. The magnitude and timing of the MicroERG peak were then measured as the average of the three most prominent channels. For spiking activity, filtering was performed in the 300-3000 Hz band. The light-evoked spike recordings were used to confirm tissue health, responsiveness, and good electrode–tissue contact.

Impedance/EIT data recorded with an active current source was processed by first performing band-pass filtering centered around the carrier frequency of the sinusoidal current source signal. In preliminary experiments oriented toward detection of fast impedance changes, a bandwidth of 1kHz was used; in further experiments, focused on detection of slow changes, we adopted a 20Hz bandwidth. For multi-channel parallel current injections, filtering was performed around each carrier frequency. Filtered data was then subject to demodulation by Hilbert Transform to extract the amplitude of the carrier signal. The resulting demodulated signals can be represented as a combination of baseline voltages (BVs), corresponding to the average voltage values in the absence of any perturbations, and voltage variations (δVs), defined as variations from baseline in response to alterations to conductivity of the tissue.

We first extracted BVs as average values of the demodulated voltages and then proceeded to compare the experimental values with theoretical BVs predicted by our model by computing a scaling factor, ideally equal to one.

The demodulated voltages were then subject again to bandpass filtering to extract the δV component and maintain only the signal frequency components associated with fast or slow impedance changes. For fast changes, only an additional high-pass filtering step was applied, resulting in a 10Hz-1 kHz bandwidth of the demodulated EIT signal. For slow changes, a 0.05-2Hz bandpass filter was applied to the demodulated signal. Coherent averaging was then applied to the δV signals to improve SNR, by averaging over the temporal triggers representing the start of a light pulse.

A temporal window of -10s to +20s centered around the trigger (t=0) was chosen as the output for each experiment where optical pulse trains were applied with 10s off-time after each pulse; in one experiment where the off-time was 5s, a -4s to +12s averaging window was chosen. The resulting averaged δVs represent a variation from baseline at the frequency of the EIT signal carrier in response to a light pattern (pulse or pulse train) and are calculated for each MEA electrode and each carrier frequency. In total, for our 32-electrodes MEA, the number of δV signals expected from each recording is 31 for single-frequency recordings and 186 for recordings performed with the six-channels parallel current source.

During EIT recordings, outliers may arise due to electrodes being damaged, fouled, or losing contact with the tissue either before or during data acquisition. To address this, the following rejection criteria were applied to exclude EIT traces, corresponding to specific combinations of measurement and current-injection pairs, from subsequent analysis:

- Traces associated with BVs>350mV in amplitude, very close to the limit of the instrument and signal saturation
- Traces from channels exhibiting drift, saturation or excessive noise in a baseline check recording performed at the beginning of the experiment
- Trace amplitude or total range >100μv
- Traces with derivative larger than 10μV/s

## IV. Results

We performed computer modelling, custom MEA manufacturing and experimental recordings to assess the feasibility of EIT-based imaging of retinal neural response to light stimulation.

### A. Modeling Results

Our pipeline for EIT image reconstruction was able to successfully reconstruct simulated local conductivity perturbations applied to our FEM model of the ex-vivo retina. Each set of simulated measurements was collected over the 6 injection pairs of our optimal protocol and 31 electrodes of the MEA; measurements were performed for 12 perturbations applied to 6 retinal layers; this resulted in a dataset of 72 measurement frames each one composed of 186 δVs.

A preliminary assessment of the simulated δV results showed that, for each perturbation, only a small subset of injection-measurement electrode combinations achieved δV values with meaningful SNR, such as SNR≥1, assuming an average recording noise of 0.1µV. Consequently, we chose to evaluate average δV magnitude on the top 10% of traces. The average δV values within this subset peaked in the RGC layer and were lowest in the IS-OS layer, which is farthest from the MEA. Mean absolute changes of top 10% δV traces were 2.01±1.31µV (RGC), 1.56±0.99µV (IPL), 0.57±0.32µV (INL), 0.30±0.16µV (OPL), 0.78±0.37µV (ONL), and 0.24±0.17µV (IS–OS). As a percentage of BV values, they were 0.129±0.072% (RGC), 0.088±0.044% (IPL), 0.015±0.007% (INL), 0.008±0.003%µV (OPL), 0.024±0.010% (ONL), and 0.008±0.002% (IS–OS).

Centre-of-Mass Error computed over the entire set of 72 perturbations was 60±31µm along the radial direction and 41±29µm along the vertical direction of the retina, corresponding to ≈13% of inter-electrode distance and ≈22% of retinal thickness, respectively.

Examples of images reconstructed from simulated perturbations are shown in Fig. 3-a) at 600µm distance from the center. A larger set of representative reconstructions in shown in Supplement Section S4. This Figure clearly shows that image reconstruction is able to correctly detect the conductivity perturbation shifting its location away from the center, but is less effective at reconstructing its shape, especially on the lower layers closer to the location of the MEA electrodes. This phenomenon is exemplified in Fig. 3-b) which shows a perturbation originally located the OPL layer. At the height of the OPL layer, approximately halfway through the mouse retina (≈100µm from MEA electrodes) the perturbation is identified with an irregular shape, as shown in third row and third column of Fig. 3.a, and in left part of Fig. 3.b. However, this perturbation is also partially reconstructed on upper layers, more distant from electrode locations, with a more regular shape. The overall trend of the radial and vertical components of the CoM error along all 72 simulated perturbations is reported in Fig. 3-c).

**Fig. 3.**
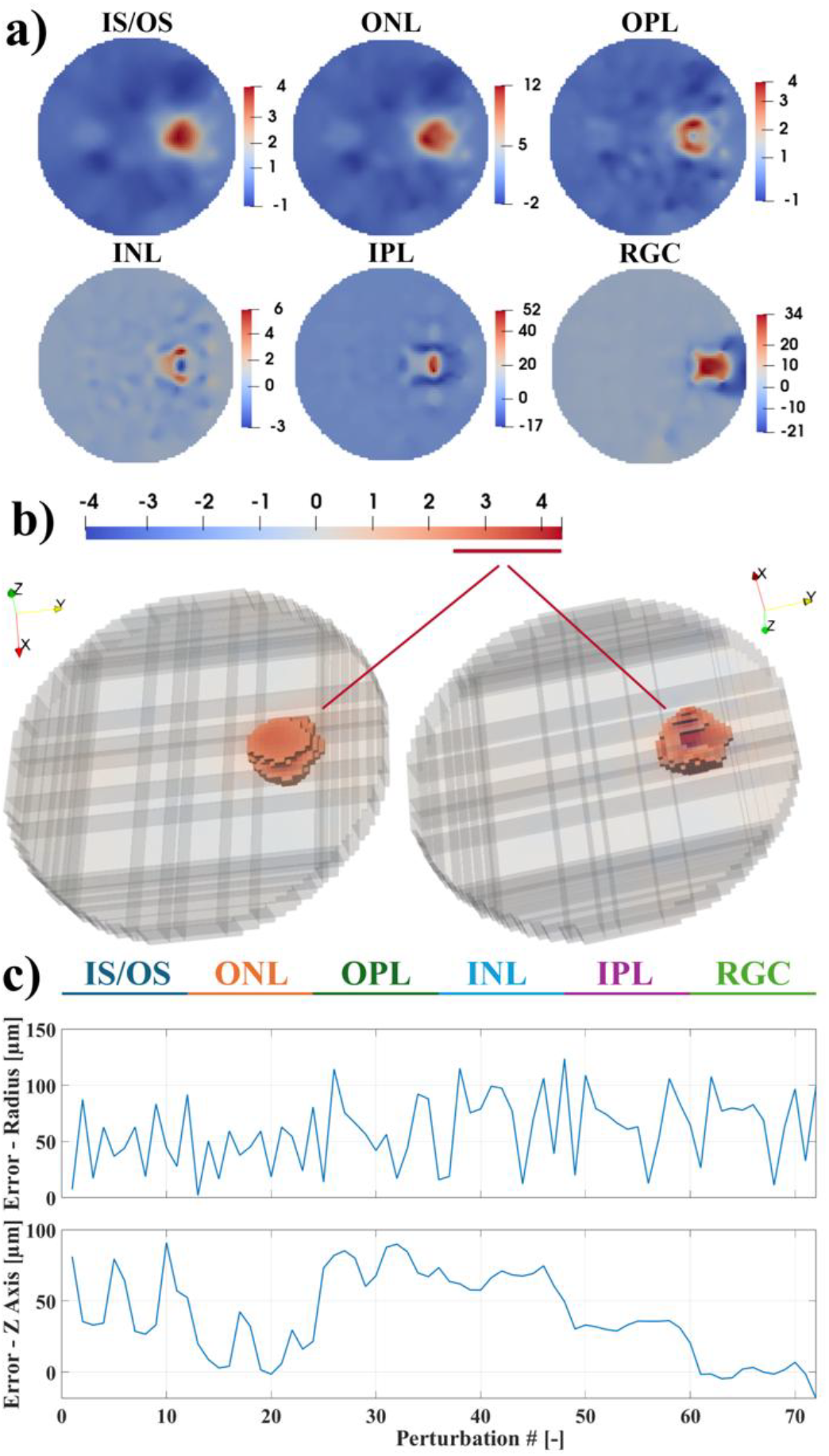
Modeling results. a) Exemplary image reconstructions from perturbations at 600µm from the center of the retinal disc, displayed at the height of their relative layer. b) Example of conductivity perturbation on the Outer Plexiform Layer (solid, threshold 50% of max value) as reconstructed over the entire retinal volume (semi-transparent). c) Error over radial and vertical axes between the Center-of-mass of applied and reconstructed conductivity perturbations. Error is shown for each layer over twelve perturbations of increasing distance from the center.

### B. Multi-Electrode Arrays Manufacturing Outcome

The Flex32 MEA we designed (Fig. 4-a-b) proved to be easy to assemble, taking ∼1h to glue 8-10 FlexPCB arrays to their support substrate and perfusion chamber. Cost of manufacture for a single MEA, when considering only consumables and chemicals involved, was approximately £30. Electrical properties of individual electrodes were in the range of 2.5 to 3.7kΩ and -32 to -17° for magnitude and phase respectively, when assessed in the experimentally relevant range of 2-10 kHz.

**Fig. 4.**
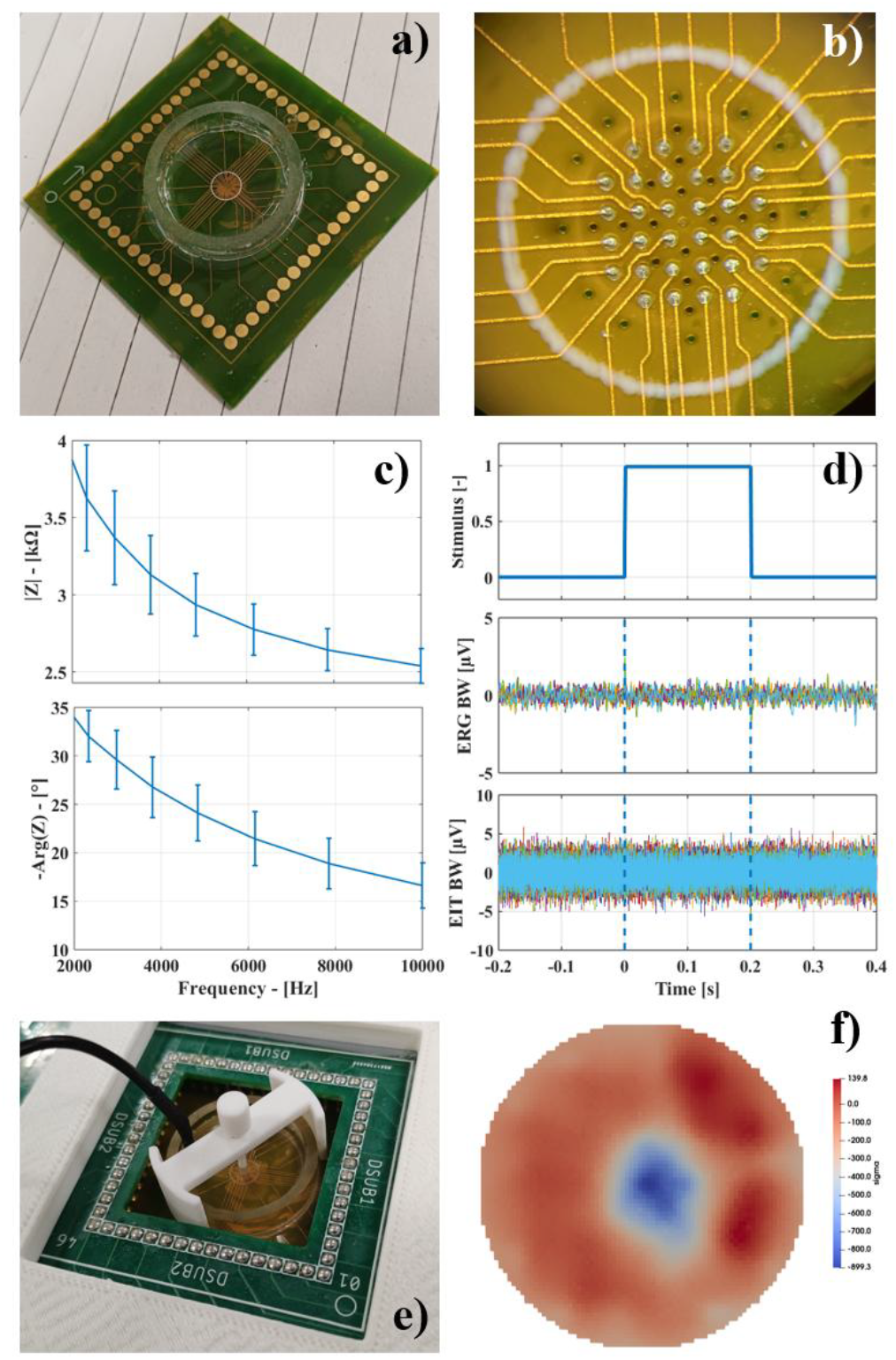
Features of the Flex32 custom MEA. a-b) Photo of the assembled MEA and zoom on platinized electrodes. C) Representative magnitude and phase of electrode impedance measured over multiple frequencies. d) Voltage readout from the MEA in the ERG and EIT/spikes bandwidths showing no sensitivity to optical pulses. e) Physical perturbation applied to the MEA to test EIT imaging. f) Image reconstruction of the test perturbation.

Compared to the commercial MEA, our Flex32 showed negligible voltage drift when recording with our neural EIT system (please see Supplement Section S1 for more information).

The Flex32 MEA we developed also showed no intrinsic sensitivity to optical pulses when tested in realistic conditions inside the experimental setup with running perfusion without any retinal tissue. An example of voltage traces recorded from the MEA electrodes before, during and after a 200ms optical pulse is reported in Fig. 4-d) and shows no traces of photoelectric artefact.

A basic test of the MEA for use in EIT showed that our combination of recording system and electrode array could successfully image a static, large perturbation placed in the middle of the imaging field (Fig. 4-e-f).

### C. Experimental Results – Standard Electrophysiology

MicroERG signals were successfully detected from N=6 retinae. MicroERG signal amplitude and occurring time of the first negative peak (a-wave) were -304.5±176.3µV (p=0.0082) and 33.0±12.2ms (p=0.0012), respectively. These values are compatible with values reported in literature for similar ex-vivo retinal recordings [38]. High-frequency spiking activity in response to optical pulses has been detected on N=1 retina; in that experiment, clear spiking response has been detected with peak amplitude of 26.1±4.7µV and 40.8±5.9ms. Figure 5 shows representative examples of MicroERG and spikes recorded from our Flex32 MEA. Supplement Section S3 reports details of an experiment in which electrophysiology was collected using our Flex32 MEA connected to a standard amplifier, rather than an EIT system.

**Fig. 5.**
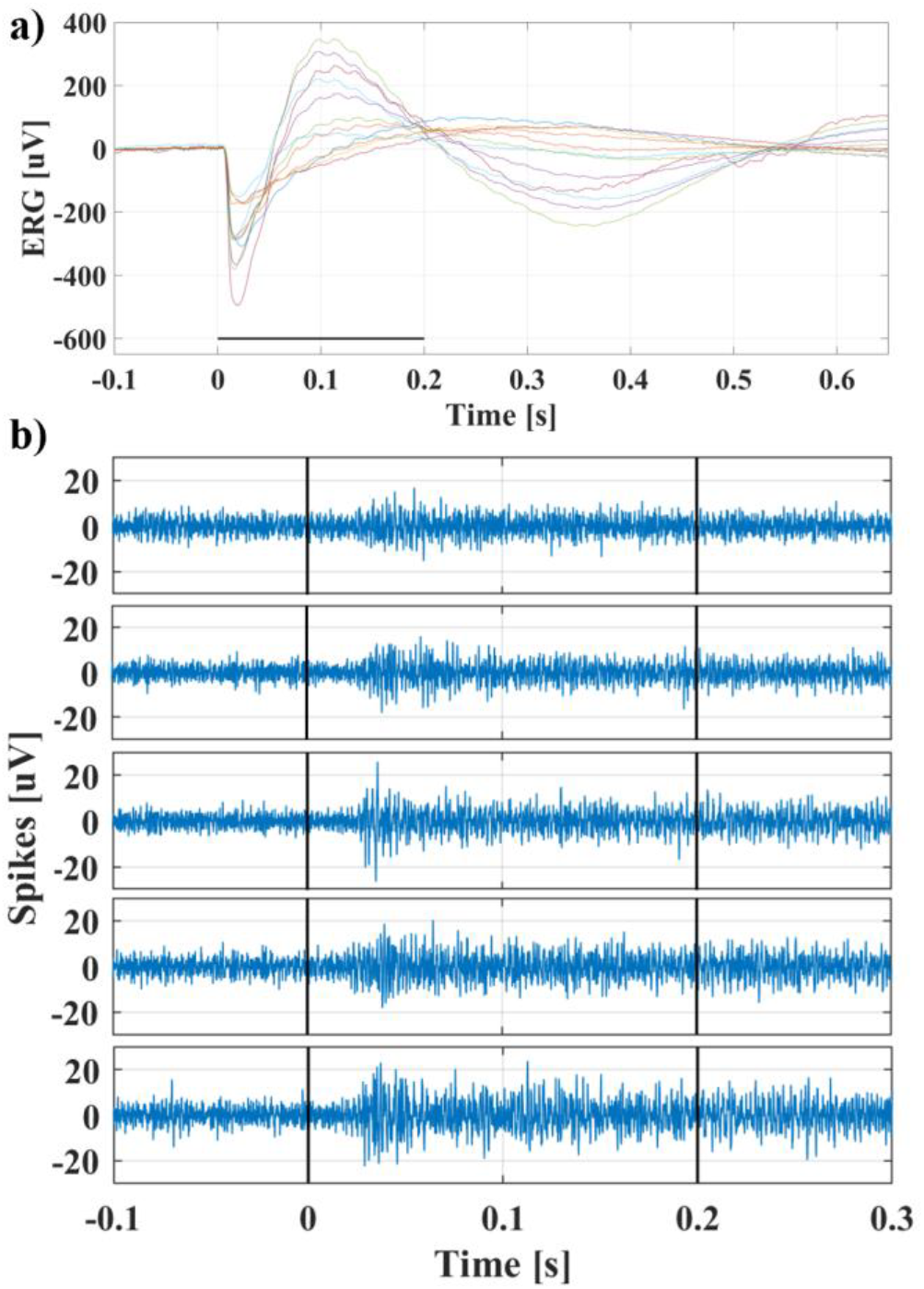
Electrophysiological markers detected from FlexPCB MEAs in response to 200ms optical pulses. Representative channels with strongest response are shown for clarity. a) MicroERG signal detected in the 1-300Hz bandwidth, b) Spiking activity detected in the 300-3000Hz bandwidth, shown for a few representative electrodes. Horizontal black line in a) and vertical black lines in b) indicate the duration of the stimulation pulse.

### D. Experimental Results –Retinal Impedance

We first compared experimentally recorded baseline voltages with those predicted by the FEM model. Since this comparison does not depend on detecting functional changes, it was based on data from all experiments, including recordings without light-evoked responses (n=27 recordings, N=5 retinae).

For each recording, we computed the scaling factor between experimental and model values for each protocol line, then took the median value across the entire recording to mitigate skewing introduced by the contact impedance of injection electrodes. The resulting scaling factor for BVs was 4.4±1.4 (p= 2.11×10^-12^vs hypothesis of scaling factor equal to one), with a range of [2.2÷7.4], indicating experimental values over 4 times higher, on average, than predicted. Since BVs are the voltages generated by the injection of fixed-amplitude current, this result indicates an overall retinal impedance significantly higher than predicted by a model built using conductivity values available from literature. Figure 6 shows the correlation between experimentally recorded BVs, and theoretical BV values predicted by the FEM model.

**Fig. 6.**
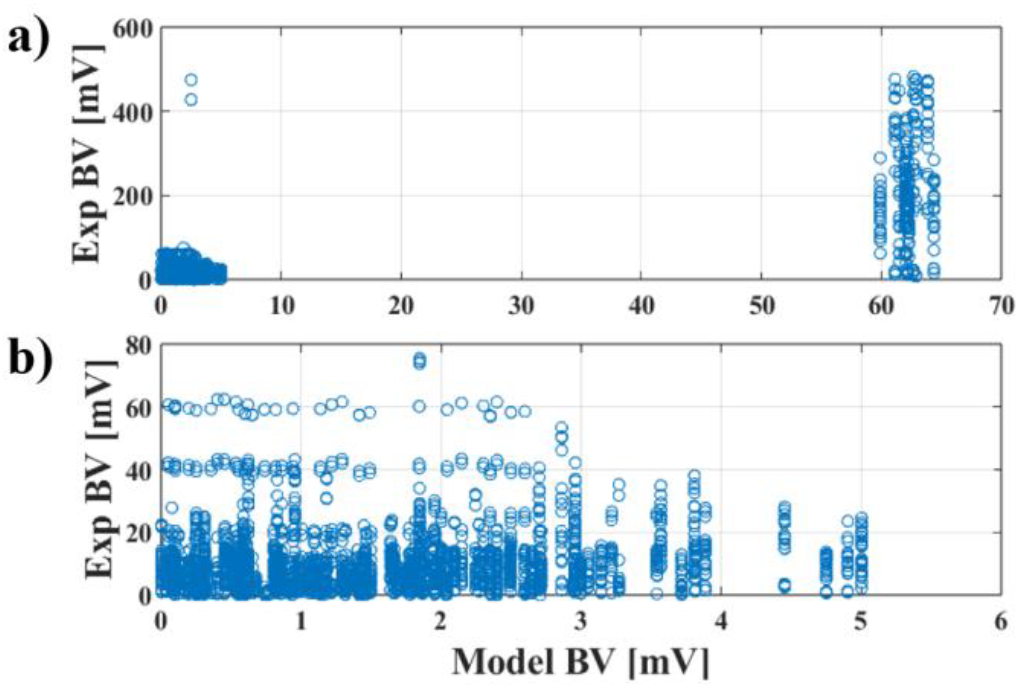
Comparison between simulated and experimental BVs. a) Full dataset. b) Subset excluding values affected by injecting-electrode contact impedance.

The second and main result from our study is the detection of slow impedance variations in response to trains of light pulses applied to the entire retina. Analysis of post-stimulus peak impedance variations in n=23 recordings from N=4 animals resulted in significant impedance variations from pre-stimulus baseline. Mean value of top 10% δV traces was 13.0±9.3µV (p=8.96×10^-7^). Mean percentage change of top 10% δV% traces, in respect to BV values, was 0.37±0.44% (p=4.78×10^-4^). Figure 7 shows examples of EIT δV data collected from each of the 4 retinae included in the dataset in response to light pulse trains. Representative δV traces recorded in response to individual light pulses, along with data illustrating signal processing within the fast neural bandwidth, are provided in Supplementary Section S6.

**Fig. 7.**
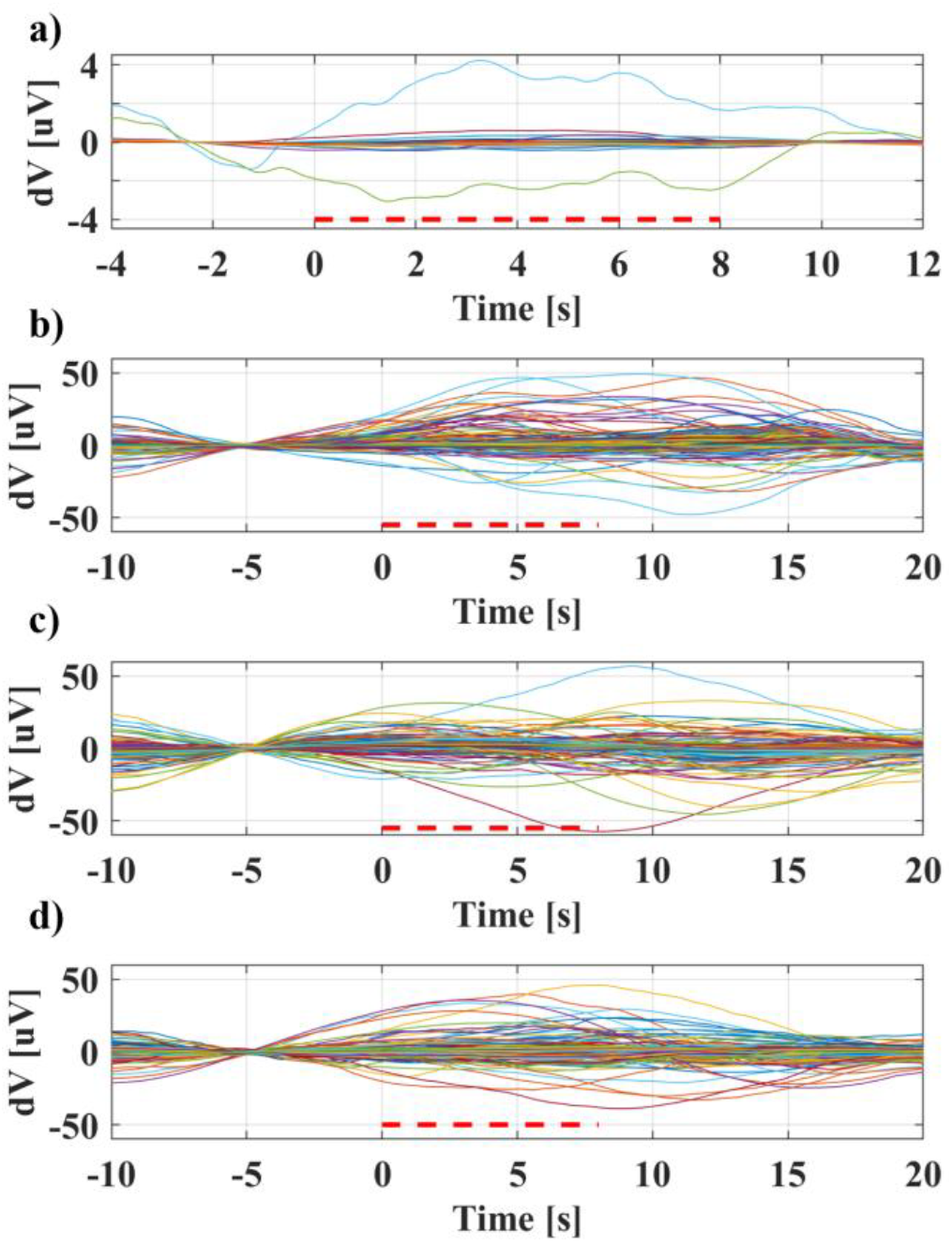
Representative voltage variations ( (δV) in response to light stimulation. a) Slow δV response to optical pulse trains (20x 200ms on/off, 5s off between bursts), measured at 6 kHz and 10µA. b) Slow δV response to optical pulse trains (20x 200ms on/off, 10s off between bursts), measured at 3.0-8.0 kHz and 10µA. c-d) Slow δV response to optical pulse trains (20x 200ms on/off, 10s off between bursts), measured at 5.2-5.7 kHz and 10µA, from two different retinae.

The third finding of our study concerns frequency dependence. All recordings were performed using sinusoidal current injection in the low-kilohertz range (2–8 kHz), applied either sequentially or in parallel. Previous investigations of neural applications of EIT have identified optimal frequencies that maximize impedance change detection for fast, action-potential–driven signals, such as approximately 1.7 kHz in the brain [5] and 6 kHz in peripheral nerves [6]. In contrast, for slower impedance changes driven by extracellular phenomena, frequency dependence has been reported to be less pronounced [9]. In this study, we evaluated the presence of frequency dependence by analysing the relationship between the normalized voltage changes δV% and the frequency at which impedance was measured. This relationship was assessed over all δV% traces collected for the study on functional impedance changes, corresponding to a total of m=2550 δV% values from n=23 recordings from N=4 retinae. The results of this analysis are presented in Figure 8 which shows no clear correlation. The correlation factor between frequency and δV% was found to be equal to R=0.079, corresponding to no correlation.

**Fig. 8.**
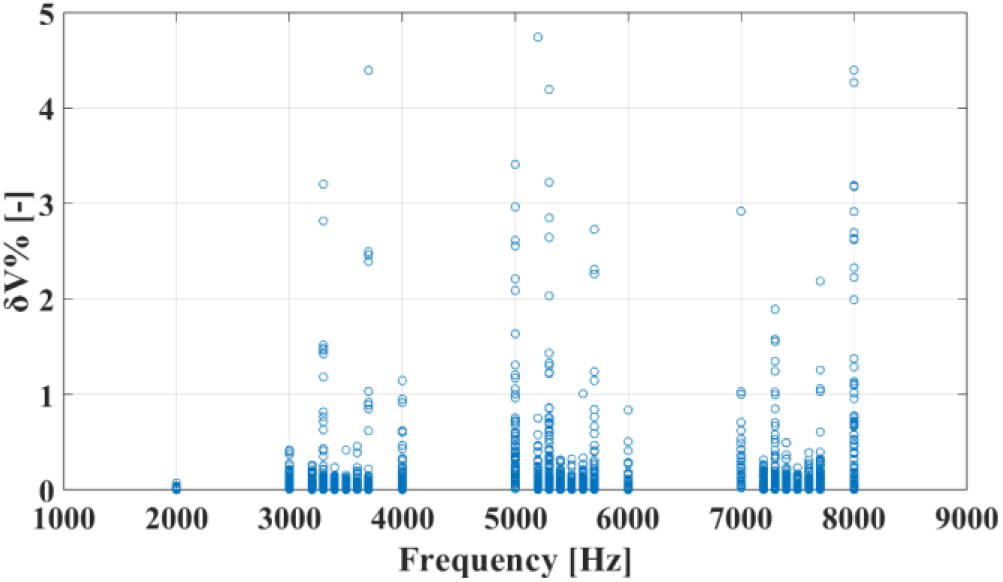
Relationship between impedance measurement frequency and relative impedance change during light stimulation (δV%).

## V. Discussion

### A. Modeling

Within this study, we first performed modeling and simulation of retinal EIT to estimate the magnitude of the expected impedance variations that we could expect from experimental recordings and the accuracy of image reconstruction. Impedance changes in the retina were modeled as discs with a diameter of 200µm and a thickness equal to that of the tissue layer in which they were located; a 5% change from baseline tissue impedance was applied. In regard to the magnitude of impedance change, previous works from our group in neural EIT imaging reported impedance changes of approximately 0.5-1% in neural tissue; we originally adopted this value for our retinal EIT modeling and later revised it to 5% based on the hypothesis that graded potentials in the first five layers of the retina could lead to larger impedance variations compared to action potentials, and to match early experimental results. However, we emphasize that this 5% value is not intended as a physiologically accurate estimate, but as a sensitivity stress-test to determine detectability limits under conservative noise assumptions. In terms of spatial geometry, we chose to model impedance changes associated with neural activation as occurring within a highly localized region, whereas physiologically retinal functional activity in response to light is not strictly confined to the area of the retinal surface directly stimulated by light or to a single layer, but rather shaped by a complex network of both vertical and lateral interactions. While we acknowledge this is a very simple approach, our main interest was not in detailed modeling of physiological dynamics but rather probe, with a reasonable test perturbation, whether retinal impedance changes were within the limits of detection of dedicated EIT hardware. Future work could focus on developing integrated models embedding both electrophysiological activity in response to stimuli and impedance changes, as done previously for peripheral nerves [39]. Our modeling results indicate that, for any given perturbation location, only a relatively small subset of injection-measurement electrode pairs exhibit significant impedance changes, likely reflecting the challenge of detecting small perturbations across a much larger MEA area. In the context of future potential retinal EIT implementations as an embedded feature of implanted retinal devices, these findings suggest that a data-compression strategy could be employed by only transmitting a selected subset of recorded voltages with large SNR to external interfaces.

Localization of perturbations using the CoM approach was achieved with a lateral error of approximately 13% of the inter-electrode distance. This performance is comparable to the localization accuracy reported in prior EIT studies of cylindrical nerve geometries [7], [40] and indicates robust lateral localization capability. Vertically, the CoM localization error was approximately 22% of the total retinal thickness, indicating that while EIT is unlikely to localize the source of an impedance change to an individual retinal layer, it may nonetheless be capable of localizing the source down to two adjacent layers.

Comparison between model BVs and experimental ones resulted in a ≈4-fold scaling factor. While such a discrepancy between theory and data is substantial, it is only a minor surprise given the large variability of conductivity values of retinal layers reported in literature. Moreover, in our model a 5% conductivity variation was applied as a localized perturbation to simulate δVs in the microVolt range, whereas neural EIT functional changes are typically modeled using 1% variations. A baseline impedance 4.4 times higher would offset this difference and require only a smaller relative change. However, this scaling factor also reflects electrode contact impedance and model simplifications, and should not be interpreted as a direct measurement of retinal tissue conductivity

### B. MEA Manufacturing and Standard Electrophysiology

Electrophysiological recordings in this study were performed using custom-made MEAs. This was done out of necessity after finding technical issues in interfacing an EIT recording system with existing commercial arrays. While the causes of this technical issue are not completely clear, we hypothesize that the DC bias compensation system of the EEG recorded embedded in our EIT setup could not handle very small, high-impedance electrodes such as the ones from the MEA used in our early tests. Future work could be performed with different commercial arrays or rely on EIT recorders with different input-stage electronics, such as AC-coupled front ends. Nonetheless, our custom MEAs provide a low-cost entry platform for impedance studies on ex-vivo tissue. The MEA we developed relies on common Flexible PCB manufacturing techniques, which has the downside of limiting electrode diameter, drill hole diameter and track spacing to ≈100µm, leading to a lower electrode count within the tissue area of 32 total electrodes, compared to 64 of the commercial MEAs. On the other hand, larger electrode size also led to a much lower contact impedance magnitude which is beneficial for bioimpedance studies. Phase angle of contact impedance for the FlexPCB MEA was in the range of ≈20-30°, which is a typical range for electrodes coated with platinum. A lower phase angle of ≈5-10° could be achieved performing coating with conductive polymers such as PEDOT, as was typical of our previous studies; however, this type of coating could not be used in the present study due to the photoelectric sensitivity we experimentally observed. Another drawback of larger electrodes is that they reduce the likelihood of detecting localized electrophysiological activity. Mice retinal ganglion cells have a size of approximately 10-20µm, which means commercial MEAs with 20µm electrodes have a much higher likelihood of detecting single-unit or few-units activity compared to our 100µm electrodes. Regardless, in a single experiment we recorded spiking activity in response to light pulses, demonstrating that in optimal experimental conditions our MEA can potentially detect this type of electrophysiological marker; more so, our main focus for this study was on recording bioimpedance variations rather than standard markers. Electrode diameter is also less meaningful for larger-scale fields such as the MicroERG, which we successfully recorded in the majority of our experiments. Future versions could be manufactured using more expensive micro-PCB manufacturing techniques which might allow smaller electrode size.

### C. Impedance Recordings and EIT applications

This study evaluated the potential of EIT as a tool for tracking functional activity in the retina. Image reconstruction of impedance changes as narrow as 200 µm was performed in simulation. In contrast, most experimental recordings employed full-field illumination across the entire tissue patch. This strategy was adopted to maximize the light-evoked retinal responses, as the primary aim of the present work in regard to the experimental components was to establish feasibility of detecting stimulus-evoked impedance changes rather than determining the minimum detectable spatial scale. Experimental measurements were also acquired using different carrier frequencies and single or parallel current sources consistently with this exploratory objective. In addition to detecting functional activity elicited by optically evoked stimuli, retinal EIT may also be applicable to tracking electrically evoked neural activity, such as that arising in retinal prosthetic systems. In fact, several prior neural EIT studies have focused on the detection of electrically evoked action potentials [7], [41], [42]. However, EIT measurements of electrically-evoked activity introduce additional challenges, most notably the need to mitigate stimulation-related artefacts [43]. Accordingly, the present study focused on responses to optical stimulation as proof of feasibility, while future work will include pulsed electrical stimulation.

The results of our preliminary experimental recordings are within an order of magnitude of the modeling predictions. The model predicted mean voltage variations of 0.24-2.01 µV, depending on the tissue layer, with an average of approximately 0.9µV or 0.05% across all simulated perturbation locations, evaluated over the top 10% traces by magnitude. Experimentally, we observed a mean δV of 13.0 µV or 0.37%, evaluated in the same conditions. Although this represents an approximately 10-fold difference between modeling and experimental results, it can be attributed to the discrepancy between the localized perturbations used in the modeling and the full-field illumination applied to the entire retinal patch ex-vivo, which likely induced impedance variations over a much larger surface area. Regarding the shape and timing of dV traces, both positive and negative voltage variations are observed, reflecting electrode proximity to current-sinking or sourcing regions, as commonly reported in the EIT literature. The apparent early onset arises from temporal broadening introduced by a narrow low-pass filter and baseline subtraction over −8 to −2 seconds pre-stimulus (zero mean at t = −5 s), with additional contribution from averaging across pulse train repetitions, where residual activity from preceding stimuli may overlap with subsequent responses. Future experiments may be performed with longer time intervals between pulse trains to ensure full return to baseline. Within the instrument’s recording bandwidth, our data did not reveal an optimal measurement frequency for detecting impedance changes. The absence of a clear frequency dependence, together with the slow temporal profile of the observed changes, parallels previously reported slow impedance variations in the brain [9] and may point to a similar extracellular physiological origin. Thus, the observed slow, stimulus-evoked impedance variations do not match typical fast neural EIT signals associated with action potentials but are likely to reflect slow extracellular and metabolic processes linked to retinal activation.

Several factors limit the strength of the experimental conclusions.

The experimental dataset we report in this study is limited in size and includes recordings acquired across a heterogeneous range of experimental parameters, which reflects the exploratory nature of the study.

While the current dataset does not clearly establish a physiological origin for the slow δVs, our control recordings exclude the most likely artefactual sources:

- the Flex32 MEA produces no photoelectric artefact under our stimulation conditions;
- the MicroERG signal disappears when the tissue is removed from the MEA, and its time course does not scale with pulse duration;
- light-evoked MicroERG and spiking signals contribute at most 0.13 µV within the EIT bandwidth, two orders of magnitude below the δVs reported here.

The experimental δVs differed from model predictions by approximately an order of magnitude, while measured BVs were about four times higher than predicted. As discussed in Section V.A, these discrepancies partly reflect the variability of published conductivity values for retinal tissue, on which the model was based, as well as the difference between the localized perturbations simulated and the full-field illumination applied experimentally. The model is therefore suitable for estimating the expected range of signal amplitudes, which guided the choice of recording hardware, averaging and current injection protocol, but not for predicting the exact value of individual measurements.

Another limitation of this study was the use of wide-field illumination, which likely activated the entire surface of the retina and prevented identification of a single localized functional activation spot. For this exploratory study, we chose to use wide-field illumination rather than localized, spot-sized retinal illumination in order to maximize the likelihood of recording functional changes.

Building on these preliminary results,, future studies will focus on confirming these observations on larger datasets recorded with a fixed protocol, pharmacological block of retinal activity, and spatially patterned stimulation.

Taken together, the modeling, technical developments, and experimental data presented in this study constitute a first step toward establishing the feasibility of EIT imaging in the retina.

## VI. Conclusions and Future Work

This study presents a comprehensive framework for the application of Electrical Impedance Tomography to ex-vivo retinal tissue, integrating computational modeling, technical methodologies, and preliminary experimental results. Our main achievements with this project were:

- Simulation-based predictions demonstrating that small, spatially localized changes in electrical conductivity can be detected as microvolt-level signal variations at individual tissue layers in ex vivo preparations.
- The development of a low-cost, custom-designed microelectrode array for retinal EIT recordings, with potential applicability to a range of ex vivo tissue patch recording studies.
- Preliminary experimental observation of slow, light-evoked impedance variations in ex vivo retina, within an order of magnitude of model predictions, requiring confirmation in larger controlled datasets.

To the best of our knowledge, this work represents the first reported experimental measurement of impedance changes acquired specifically for EIT-focused retinal imaging. Collectively, these results indicate the feasibility of retinal EIT and provide a foundation for further methodological and technological development. Future work will focus on advancing MEA and recording system performance, expanding data acquisition across a broader range of experimental conditions, such as including more complex visual stimulation paradigms, and extending retinal EIT toward in vivo applications.

## Data Availability

Modeling code and experimental data are available on the UCL Research Data Repository: https://rdr.ucl.ac.uk/. Electronic designs, such as those of the EIT current sources used for this project, are available on the UCL EIT group GitHub page: https://github.com/EIT-team.

