## Supplemental methods and data for "Toward Electrical Impedance Tomography of Functional Activity in the Retina"

### Summary of Content

This document provides further information and data for the manuscript "Toward Electrical Impedance Tomography of Functional Activity in the Retina". The document contains the following sections:

S1: Comparison of electrical baseline drift in our early electrophysiological recordings collected with commercial MEAs against the ones performed with our Flex32 MEA.

S2: Details of an intermediate-stage MEA design, initially developed for use in this study, but ultimately discarded due to the identification of significant photosensitivity artifacts.

S3: Control experiment where our custom Flex32 MEA is connected to a standard commercial electrophysiology amplifier, rather than an EIT recording setup.

S4: Larger set of representative image reconstruction results from simulated perturbations.

S5: Overview of the experiments included in our dataset.

S6: Control recordings and data analysis.

### 1. Electrical Drift in MEA Recordings

We originally planned to perform electrophysiological and EIT recordings using a combination of our existing neural EIT system (ScouseTom) and commercially available MEAs.

As explained in Section III.B of the main document, our MEA of choice was the 60PEDOT-pMEA200/30iR-Ti-Au (Multi Channel Systems, Reutlingen, Germany). This MEA featured 60 electrodes of 30µm diameter arranged in an 8x8 grid configuration with missing corners, intercalated with through-holes of diameter 20-90µm which allowed suction to improve tissue adherence to the array. Electrodes were made of Titanium-Gold alloy and coated with PEDOT-CNT. This MEA was chosen on the basis of previous successful use by collaborators in retinal electrophysiology experiments in combination with a standard electrophysiology amplifier Intan 512 Ch RHD recording system (Intan Technologies; Los Angeles, CA). However, standard electrophysiology amplifiers usually feature an analog-to-digital resolution of 16 bits and a sampling rate in the order of 20-30kHz, which was not viable for this study, as past neural EIT studies demonstrated the need for higher sampling rates and vertical resolution.

For this study, we initially decided to connect our existing EIT setup to the commercial MEA from MultiChannel Systems (MCS-MEA for brevity) though a custom PCB adapter to perform Ephys/EIT recordings. The recording module in our EIT system is a commercial high-resolution, high-speed EEG/biopotential amplifier (Actichamp, Brain Products GmbH, Germany) with 24-bit resolution, 10kHz bandwidth and 50khz sampling rate. However, we quickly identified the presence of a significant baseline drift in our recordings which quickly saturated voltage on the amplifier channels and blocked us from performing sufficiently long recordings with this setup.

This issue was replicated on different preliminary experiments and was even present on saline-filled MEA with no retinae, indicating it was not tissue-related. Switching reference electrodes in the MEA bath also did not remove the drift. Our attempts to diagnose the source of this technical issue and to solve it, for example by adding an interconnecting stage between the MEA and EEG device to perform either AC-coupling or DC-biasing, were ultimately unsuccessful due to the closed-source nature of the amplifier. However, because this baseline drift did not occur with AC-coupled systems like the Intan, we hypothesize that it arises from a spurious interaction between the MEA electrodes and the DC-coupled ActiChamp input-biasing circuitry, leading to charge accumulation on the MEA electrodes. Ultimately, since the use of the existing biopotential recording system was a necessity for our study due to the technical demands of recording EIT signals, we decided to proceed with the development of the Flex32 MEA described in the main document.

Overall, switching from the commercial MCS to our Flex MEA resulted in a functionally complete removal of the baseline drift (1.25±0.62 vs 0.22±0.10, p=0.023, n=5) and justified adoption of the new MEA for this study. Supplementary Figure 1 shows a baseline comparison from a

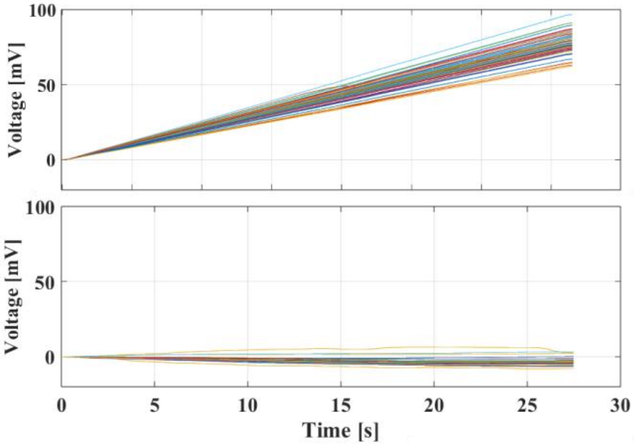


Fig. 1. Voltage baseline drift in original MCS MEA (top) and custom Flex32 MEA (bottom).

recording performed with a mouse retina mounted on the MCS MEA and connected to our recording system, with no light stimulation applied. The signals were low-pass filtered at 1 Hz for this plot to improve visibility of the drift.

### 2. Photoelectrical Artefact in PEDOT-coated MEAs

The decision to abandon commercial MEAs in favor of a custom design on the basis of persistent electrical drift led us to consider multiple manufacturing and coating technologies.

For MEA manufacturing we settled on traditional PCB technology, consisting of copper traces on FR4 substrate with exposed MEA pads and ENIG surface finish. To reduce magnitude and phase of contact impedance between electrode and tissue, and thus improve outcome of our impedance measurements, additional surface coating was required.

Our first material of choice was PEDOT:pTS, a conductive polymer frequently used for this purpose which offers significant reduction in contact impedance compared to uncoated metal electrodes, and which our group employed for several previous studies (See References in main manuscript). However, in a limited set of experiments using PCBs with PEDOT-coated electrodes, we observed substantial photoelectrical artefacts in the light-evoked retinal recordings obtained with this type of MEA.

Supplementary Figure 2-a shows an example of PEDOT-coated PCB MEA and the electrical response recorded from the array placed on the perfusion system with no retina mounted. Periodic light stimulation pulses of 500ms width were applied to the MEA. Top panel of Fig. 2-b shows the lack of meaningful signals when light pulses are present but the pathway to the MEA electrodes is physically obstructed. Fig. 2-b middle and bottom panels show how an artefactual signal is present when light pulses are allowed to illuminate the MEA electrodes, and how the artefact increases in amplitude when switching from yellow to blue light.

The absence of retinae in this test confirms that the observed signal changes are artefactual. Additionally, the lack of changes when the light pathway is obstructed further supports our hypothesis that the recording artefact is directly attributable to light exposure on the array, rather than to electromagnetic interference within the experimental setup.

Photoelectrical artefacts are a known and previously reported effect in neural engineering Kozai and Vazquez 2015, see Refs in main document), however, to the best of our knowledge there is no prior data explicitly showing the effect of light on PEDOT-coating PCB copper/gold traces. In the end, the strong photosensitivity observed in PEDOT-coated PCB electrodes prompted us to seek an alternative coating method, leading to the adoption of platinum coating described in the main document. As shown in Fig. 4d of the main document, platinization of the Flex32 MEAs reduces the contact impedance to suitable levels for EIT measurements without exhibiting any detectable photosensitivity.

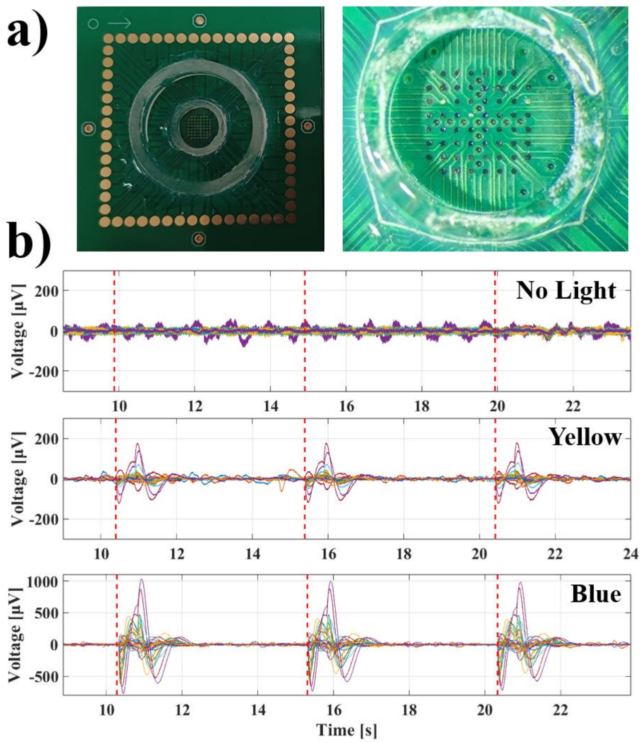


Fig. 2. A) PCB-based MEA and detail of PEDOT-coated electrodes. B) Voltage recorded in ERG band with no mounted retina under no light stimulation (top), yellow light stimulation (middle) and blue light stimulation (bottom).

### 3. Performance of Custom MEA with AC-coupled Recording Systems

As explained in the main document and in this supplement, our custom Flex32 MEA was designed and manufactured to ensure drift-free recordings when connected to our existing Fast Neural EIT system. However, before performing experiments with the combination of the Flex32 MEA and Actichamp-based EIT system, we performed one control experiment in which the novel array was connected to a standard Intan 512 Ch RHD recording system (Intan Technologies; Los Angeles, CA) commonly used for standard retinal electrophysiology. The purpose of this control was to confirm the capability of the new MEA to record standard electrophysiology markers such as MicroERG and spiking activity before moving to the collection of novel impedance-based data with the EIT system. We only performed a control experiment rather than a complete one because this amplifier was designed for common electrophysiology signals and had a limited input voltage range which was not compatible with impedance-based recordings (4mV vs 400mV of our neural EIT system). The new custom MEA was able to successfully detect MicroERG signals and retinal spikes. Over n=5 recordings, average MicroERG signal peak amplitude and time were -133±43μV ($p= 0.001$) and 48±12ms ($p= 0.002$), respectively, and spiking response peak amplitude and time were 8.17±0.94μV ($p=4.2370e-05$) and 66±15ms ($p=6.34e-04$), respectively. These values are compatible with values reported in the main document and in literature for similar ex-vivo retinal recordings and confirm the feasibility of using our novel

MEA for electrophysiological recordings. Supplemental Figure 3 shows representative examples of MicroERG and spikes recorded within this control experiment in response to optical pulses.

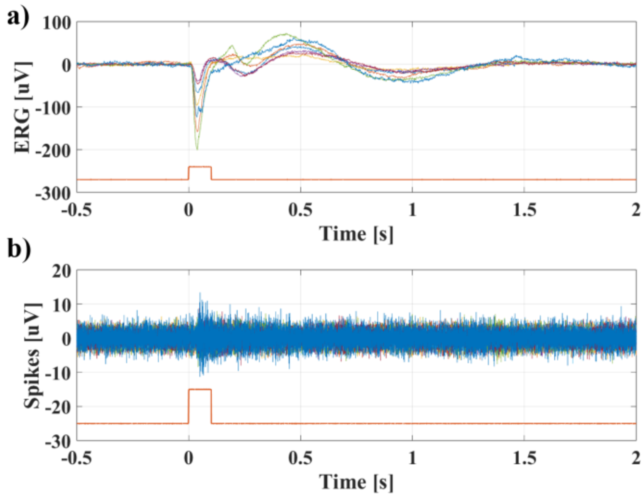


Fig. 3. Examples of electrophysiological markers detected from FlexPCB MEAs in response to optical pulses. a) MicroERG, b) Spiking activity.

### 4. Results of Image Reconstruction from Simulated Data

Examples of images reconstructed from simulated perturbations are shown in Supplemental Fig. 4, at regular distance intervals of 0-300-600-900µA from the center.

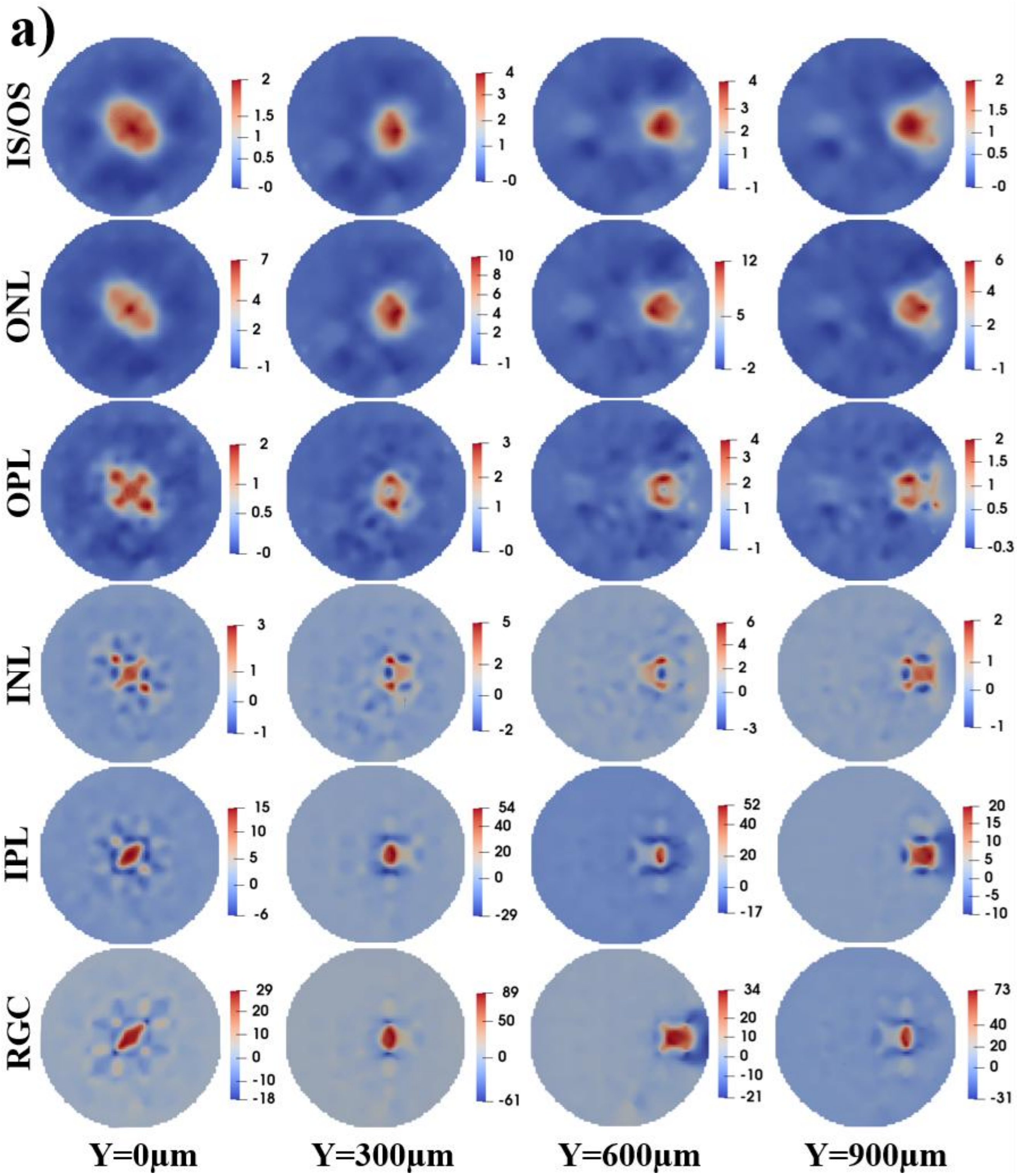


Fig. 4. Representative image reconstructions from perturbations at 0, 300, 600 and 900µm from the center of the retinal disc, displayed at the height of their relative layer.

### 5. Overview of Experimental Dataset

Within this study, experiments were performed on retinae explanted from a total of 26 mice. Out of these, 16 experiments were dedicated to technical development and were affected by drift and photoelectric sensitivity MEA issues as described in Sections S1 and S2 of this Supplement. The remaining 10 experiments were performed using the final FlexPCB version of the MEA. Of these remaining 10 experiments, 3 experiments were excluded from the dataset due to the absence of light-evoked electrophysiological markers, consistent with either reduced tissue viability or suboptimal electrode–tissue interface. The remaining 7 experiments constitute the core dataset for our study and are listed in Supplementary Table S-I. More specifically, out of these 7 experiments:

- 1 experiment was dedicated to testing the novel MEA with an established electrophysiology setup, which did not allow EIT/impedance recordings; the results are reported in Supplement Section 3.
- 1 experiment allowed successful recording of both MicroERG and retinal spikes, but impedance data was not recorded successfully due to a battery malfunction in the current injection circuitry. Data from this experiment is included in MicroERG statistics and an example of spike recording is shown in the main document.
- 1 experiment allowed successful recording of MicroERG and featured impedance data collection at single individual frequencies in response to individual periodic light pulses. N=4 experiments featured successful MicroERG recording, and single- or multi-frequency data collection in response to optical pulse trains. δV markers in the main document are extracted from these data.

### 6. Control Recordings

A series of control recordings were performed to confirm that the collected data were not of artefactual origin.

The following controls were performed to demonstrate that data collected with the final FlexPCB MEA design where not the product of photoelectrical artefacts:

- <u>Pulse Duration</u>: Light pulses were applied to the same retina with durations of either 200 ms or 10 ms. As shown in Figure 5-a), the MicroERG signal exhibited nearly identical temporal dynamics across both pulse durations, suggesting that the signal is a true biopotential rather than an artifact, which would be expected to scale in timing with pulse duration.
- <u>Presence or Absence of Retinal Tissue</u>: Light-evoked MicroERG recordings were first performed with retinal tissue mounted on the MEA and then repeated after the tissue was removed at the end of the experiment. As shown in Figure 5-b), the MicroERG signal disappeared, whereas a photoelectric artifact would have persisted.

Another set of controls was carried out to demonstrate that standard electrophysiological signals such as MicroERG and spiking, generated in response to light pulse, do not have significant signal components in the frequency band used in this work to perform impedance measurements. Figures 5-c) and 5-d) show respectively how two signals with visually observable light-evoked MicroERG and spiking activity appear when filtered with a 20Hz passing band centered around carrier frequencies ranging from 1 to 8 kHz in 1 kHz steps. During peak MicroERG, signal values in the EIT bandwidth reached a mean value of 0.04±0.03μV across all carrier frequencies and an overall maximum of 0.08μV, compatible with background fluctuations outside of peak electrophysiological activity, and much lower than the actual light-evoked impedance variations reported in the previous section. For the recording with spiking activity, mean and maximum signal values in the EIT bandwidth during spiking reached similar values of 0.04±0.04μV and 0.13μV, respectively.

As described in the main text, our analysis focused on the effects of light pulse trains on the slow components of impedance variations, as neither individual light pulses nor higher-frequency signal processing yielded meaningful results. For reference, single-frequency measurements obtained from the same experiment under single-frequency acquisition mode are shown in Fig. 6-a. Representative δV traces recorded in response to individual light pulses, together with data illustrating processing within the fast neural bandwidth, are presented for comparison in Fig. 6-b) and 6-c).

TABLE S-I
Experimental Dataset

| Exp # | Light Pulses | MicroErg | Spiking Activity | EIT / Impedance Measurement |
|---|---|---|---|---|
| 1 | Single bursts of 20x pulses, 200ms on/off | Yes | Yes | - |
| 2 | 20x consecutive 200ms pulses, 5s rest between bursts | Yes | No | Single Injection Channel - Frequency: 2 to 8khz |
| 3 | 20x consecutive 200ms pulses, 10s rest between bursts | Yes | Yes | - |
| 4 | 20x consecutive 200ms pulses, 10s rest between bursts | Yes | No | Six parallel injection channels – 3.0-8.0kHz |
| 5 | 20x consecutive 200ms pulses, 10s rest between bursts | Yes | No | Six parallel injection channels - Multiple recording at low (3.2-3.7kHz), mid (5.2-5.7kHz) or high (7.2-7.7 kHz) frequencies |
| 6 | 20x consecutive 200ms pulses, 10s rest between bursts | Yes | No | Six parallel injection channels - Multiple recording at low (3.2-3.7kHz), mid (5.2-5.7kHz) or high (7.2-7.7 kHz) frequencies |
| 7 | Single 200ms pulse every 5s | Yes | No | Single Frequency: 2 to 7khz |

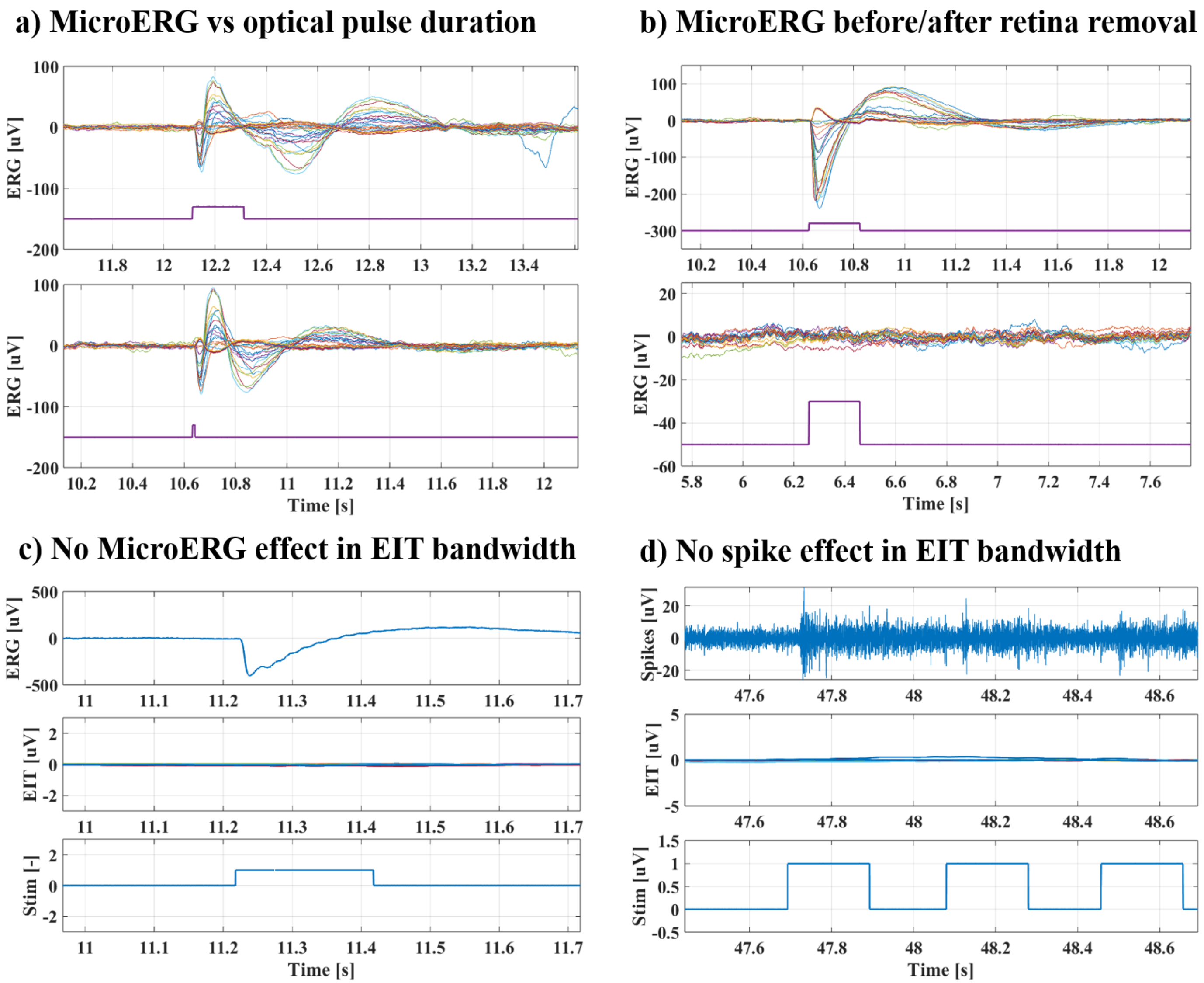


Fig. 5. Electrophysiology Controls. a) MicroERG recording from the same retina in response to 200ms and 10ms light pulses. b) MicroERG recording from the same experiment before and after removing the retina from the MEA. c) Example of MicroERG signal recorded from a single channel (top), in response to optical pulses (bottom), with no on-going current injection, and the result of filtering the signal into EIT bandwidth (mid). d) Example of retinal spiking signal recorded from a single channel (top), in response to optical pulses (bottom), with no on-going current injection, and the result of filtering the signal into EIT bandwidth (mid).

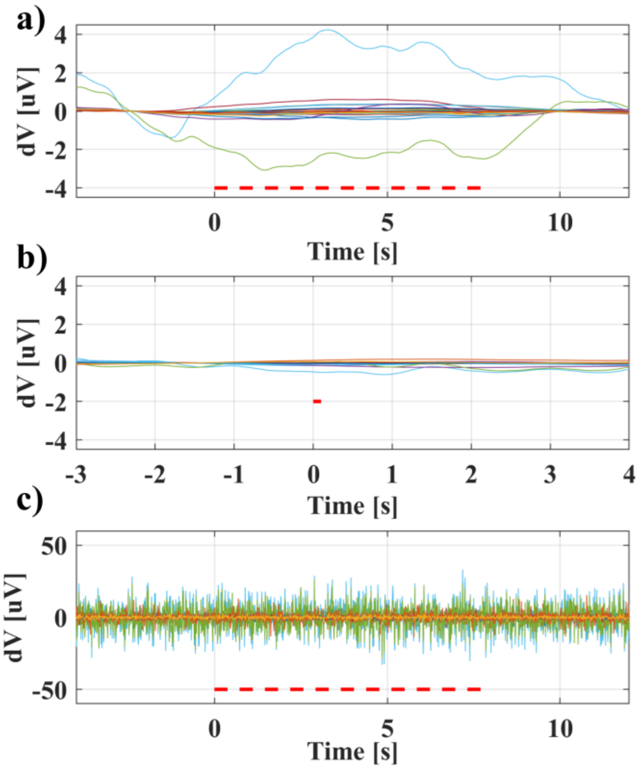


Fig. 6. a) Slow δV response to optical pulse trains (20x 200ms on/off, 5s off between bursts), measured at 6 kHz and 10μA. b) Slow δV signal response to single light pulses (1x200ms on/off, 5s repetition), measured at 6 kHz and 10μA, from the same retina as plot a). c) Fast Neural analysis outcome from same retina and recording as a), showing no meaningful response to light in this bandwidth.
